# Influence of Memory Modulation on Affective States in Methamphetamine Use Disorder

**DOI:** 10.64898/2026.11.13.766209

**Authors:** Sarah I. Qundes, Jennifer L. Stewart, Maria Ironside, Martin P. Paulus, Maëlle C.M. Gueguen

## Abstract

**Background:** Negative affect and low positive affect are linked to craving and relapse in methamphetamine use disorder (MUD). Brief positive autobiographical memory recall improves mood in non-clinical samples, but it hasn’t been tested in people in treatment for MUD. We tested whether positive memory recall improves affective state, withdrawal and craving compared with a neutral recall condition.

**Methods:** Adults with past-year MUD in their first year of abstinence were recruited from two residential treatment programs in Tulsa, Oklahoma. Participants were randomized, blinded to condition, to recall personally meaningful positive memories (Positive; n = 37) or emotionally neutral procedural memories (Procedural; n = 36) during a laboratory session. Positive and negative affect (PANAS, the prespecified primary outcomes), withdrawal (AWQ), craving (visual digital scale; Stimulant Craving Questionnaire), and anhedonia (SHAPS) were assessed before and approximately one hour after the recall. Group × Time effects were tested with mixed ANCOVAs adjusted for sex and age, with Holm and false discovery rate correction. Baseline-adjusted end-of-session differences were estimated for each outcome.

**Results:** Of 80 randomized participants, 73 began the session and were analyzed. The Group × Time interaction was not significant for positive affect (p = .689) or negative affect (p = .399). Baseline-adjusted end-of-session differences were small and imprecise (positive affect: −0.53 points, 95% CI −2.60 to 1.53; negative affect: 0.54 points, 95% CI −0.90 to 1.99). Across both groups, negative affect, withdrawal, craving and desire to use decreased over the session, while positive affect also declined (all p ≤ .015). No secondary or exploratory outcome differed between conditions.

**Conclusions:** A single positive autobiographical memory recall did not produce a detectable end-of-session difference in affect, withdrawal or craving relative to neutral recall in individuals with MUD. Affective and craving states shifted across the session regardless of memory content. Future work should test repeated or more intensive positive-affect interventions in this population.

## Introduction

Negative affect is a prominent feature of early abstinence from methamphetamine. Through hyperkatifeia, a heightened negative emotional and motivational state, negative affect drives craving and continued drug-seeking as a form of negative reinforcement (Koob, 2021; Koob & Volkow, 2009), and elevated negative affect during this period has likewise been linked to increased withdrawal symptoms and relapse risk (Wang et al., 2013; May et al., 2020). Because this negative affective state is theorized to drive continued drug use, mood itself represents a potentially modifiable treatment target.

The clinical stakes of this early withdrawal period are considerable. Methamphetamine use disorder (MUD) is characterized by a chronic, relapsing course, with an estimated 36% of individuals who use methamphetamine relapsing within six months of treatment and 61% relapsing within one year (Brecht & Herbeck, 2014). Because no medication is currently approved by the U.S. Food and Drug Administration specifically for methamphetamine use disorder, psychosocial interventions remain the primary evidence-based treatment option, underscoring the need to identify modifiable psychological targets, such as affective state during early abstinence, that could be feasibly integrated into behavioral treatment (Brecht & Herbeck, 2014).

Consistent with this model, affect itself appears to be a key mechanism of behavior change in psychosocial treatment: elevated positive affect has been shown to predict better substance use treatment outcomes and greater distress tolerance, independent of baseline negative affect (Swan et al., 2020). Existing interventions that directly target positive affect include behavioral activation, savoring- and gratitude-based exercises, and structured positive affect treatment protocols, several of which have shown promise for improving affect and coping in substance-using populations.

A complementary perspective is offered by the broaden-and-build theory of positive emotions, which proposes that positive affect momentarily broadens an individual’s attention and thought– action repertoire, thereby building durable psychological, social, and coping resources over time (Fredrickson, 2001; Fredrickson & Losada, 2005). Applied to addiction, this framework suggests that positive affect may sensitize individuals to non-drug sources of reward and support the sustained use of cognitive-behavioral and social coping strategies needed to maintain abstinence (Ashby et al., 1999; Carrico et al., 2013). Supporting this account, a study of treatment-seeking individuals with methamphetamine misuse found that higher positive affect, independent of negative affect, was associated with greater approach-oriented coping and self-efficacy for managing methamphetamine triggers, both of which are theoretically linked to reduced substance use (Carrico et al., 2013).

Autobiographical memory recall represents one promising, low-burden avenue for modulating affective state in this population. Deliberately recalling positive personal memories has been shown to reliably increase momentary positive mood relative to recalling neutral or negative memories in individuals with alcohol use disorder (Nandrino et al., 2017). Because these effects have primarily been demonstrated in non-clinical or single-disorder samples, direct testing of brief positive memory interventions within treatment-seeking clinical populations is needed rather than assuming their benefits generalize.

While many withdrawal symptoms resolve within the first week of abstinence, dysphoric mood and craving remain elevated much longer, with craving persisting for at least five weeks (Zorick et al., 2010) and prospectively predicting methamphetamine use within one to seven weeks of assessment (Galloway et al., 2008); allostatic models of addiction propose that this negative affect can persist well beyond the acute phase as part of a protracted withdrawal syndrome, in which subthreshold dysphoria, anhedonia, and stress reactivity continue to elevate relapse risk for months into abstinence (Koob, 2021). Together, these findings indicate that affective disturbance in MUD follows a protracted course extending well beyond acute withdrawal-supporting an examination of affect across the first year of abstinence rather than the acute withdrawal window alone-and that mood itself is a modifiable, low-burden treatment target whose responsiveness to brief positive memory recall has not yet been tested in treatment-seeking clinical populations. The present study examined whether a brief positive autobiographical memory-based mood manipulation could improve affective state in treatment-seeking individuals with MUD. Based on prior evidence that positive affect manipulations can reduce distress and improve psychological functioning in varying substance-using populations (Carrico et al., 2013; Stone, 2023), we hypothesized that participants assigned to the positive memory recall condition would demonstrate greater increases in positive affect and greater reductions in negative affect, withdrawal symptoms, and craving compared to participants assigned to a procedural memory recall condition. This report is limited to self-reported affective, withdrawal, and craving outcomes collected before and after this manipulation; concurrently collected neuroimaging and in-scanner cognitive task data (from the same MRI session) are not analyzed here and will be reported separately.

## Methods

### Participants

Participants were treatment-enrolled adults with MUD recruited from GRAND Addiction Recovery Center and Women in Recovery, residential substance use treatment programs in Tulsa, Oklahoma. Eligible participants were between 18 and 65 years of age, English proficient, met DSM-5 criteria for past-year MUD, and were in their first year of abstinence. Abstinence duration ranged from 21 to 355 days (median = 169 days).

Exclusion criteria included severe traumatic brain injury (Tulsa Head Injury Screen score ≥ 3), medical conditions that would interfere with study participation, including major surgery requiring general anesthesia within 12 weeks before screening, incomplete recovery from surgery, planned surgery during the study period, significant cognitive impairment, or clinically relevant or progressive medical conditions (e.g., liver, kidney, cardiovascular, respiratory, vascular, brain, metabolic, or thyroid disorders) that could affect the course of the study.

Medical, psychiatric, or other conditions that restricted an individual’s ability to interpret study information, provide informed consent, adhere to study procedures, or complete the study were also exclusionary. Additional exclusion criteria included current psychotic disorders, Bipolar I disorder, or major depressive disorder with psychotic features as determined by the Mini International Neuropsychiatric Interview (MINI; Sheehan et al., 1998), MRI contraindications, positive alcohol breathalyzer results, or positive urine drug screens (with the exception of cannabis and prescribed benzodiazepines). Individuals who were unable to comply with study procedures were also excluded. Participants were enrolled between July 2024 and March 2026. Of 273 individuals assessed for eligibility, 193 were excluded before randomization: 70 for a psychotic disorder, bipolar disorder, or current psychotic symptoms; 49 for a primary drug other than methamphetamine; 17 because they were discharged, left the treatment program, or could not be scheduled; 16 for an MRI contraindication; 15 for other medical or neurological exclusions (e.g., traumatic brain injury, seizure history, pregnancy); 14 because they were at or approaching one year of abstinence; 6 for a body mass index outside the Core protocol range; 1 who declined; and 5 for other reasons (Supplemental Figure S2). A total of 80 eligible individuals were randomized, 40 to each condition. Six participants withdrew after randomization (three assigned to the Procedural condition and three assigned to the Positive condition; reasons included leaving the recruitment site’s treatment program or no longer wishing to participate) and one participant randomized to the Procedural condition was subsequently excluded because a court-ordered ankle monitor (an MRI contraindication) was identified. None of these seven participants completed the pre-session assessment, so no outcome data were collected for them; the analytic sample therefore comprises all randomized participants who began the study session, resulting in a final analytic sample of 73 participants (Procedural: *n* = 36; Positive: *n* = 37; **Table 1**).

**Table 1.** Sample Demographics and Clinical Markers.

| **Characteristic** | **Procedural (n = 36)** | **Positive (n = 37)** | **Test** | **p** |
|---|---|---|---|---|
| Age, years (M ± SD) | 38.83 ± 9.02 | 39.14 ± 9.68 | t<sub>71</sub> = −0.14 | 0.891 |
| Female, n (%) | 28 (77.8) | 31 (83.8) | χ<sup>2</sup> = 0.425 | 0.515 |
| Abstinence duration, days (M ± SD) | 151.7 ± 112.1 | 159.5 ± 113.7 | W = 1322 | 0.917 |
| Race, n (%) |  |  | χ² = 6.30 | 0.178 |
| American Indian | 6 (16.7) | 3 (8.1) |  |  |
| Asian | 1 (2.8) | 0 (0.0) |  |  |
| Black/African American | 1 (2.8) | 3 (8.1) |  |  |
| Multiracial | 9 (25.0) | 4 (10.8) |  |  |
| White | 19 (52.8) | 27 (73.0) |  |  |
| CTQ Total Score (M ± SD) | 58.75 ± 23.33 | 55.14 ± 22.93 | t<sub>71</sub> = 0.67 | 0.506 |
| PROMIS Anxiety (M ± SD) | 53.59 ± 9.86 | 52.22 ± 9.81 | t<sub>71</sub> = 0.59 | 0.554 |
| PROMIS Depression (M ± SD) | 57.85 ± 10.21 | 56.59 ± 9.93 | t<sub>71</sub> = 0.53 | 0.595 |
| Comorbid Diagnoses†, n (%) |  |  |  |  |
| Cannabis use disorder | 22 (68.8) | 17 (48.6) | Fisher's exact | 0.137 |
| Major depressive disorder | 18 (56.2) | 15 (42.9) | Fisher's exact | 0.332 |
| Opioid use disorder | 14 (43.8) | 10 (28.6) | Fisher's exact | 0.214 |
| Alcohol use disorder | 11 (34.4) | 13 (37.1) | Fisher's exact | 1.000 |
| PTSD | 11 (34.4) | 3 (8.6) | Fisher's exact | 0.015 |
| Antisocial personality disorder | 11 (34.4) | 6 (17.1) | Fisher's exact | 0.160 |

*Note. †Comorbid diagnoses reflect a structured diagnostic interview; data were unavailable for 4 Procedural and 2 Positive participants (comorbid-diagnosis percentages are of n = 32 Procedural and n = 35 Positive participants with interview data). Group comparisons used Fisher’s exact test given small expected cell counts.*

### Study Design

Participants were randomly assigned to either a positive mood manipulation (Positive group) or a procedural mood manipulation (Procedural group). Random assignment used block randomization with a block size of ten to maintain balanced group sizes as enrollment progressed; the randomization sequence was generated and held by the study’s principal investigator and provided directly to the research assistant conducting each session. The research assistant remained blind to condition assignment until the participant entered the MRI scanner, at which point unblinding was necessary to identify and administer the correct memories for the mood-manipulation procedure; pre-manipulation assessments were therefore collected under blinded conditions, whereas post-manipulation assessments were administered by an unblinded research assistant. Participants were blind throughout the study to the existence of two conditions and to their own group assignment and were informed only that they would be recalling personal memories. The study protocol (LIBR #2024-002; version dated March 19, 2024) specified the design, primary outcomes, and primary analysis, and the trial was registered at ClinicalTrials.gov (NCT06410196; first submitted May 8, 2024) prospectively, before the first participant was enrolled (July 26, 2024). The registered outcome measures were not modified in any later version of the registration. The protocol planned enrollment of 80 participants (40 per condition) to yield 72 completers, assuming 10% attrition; no formal power calculation was reported. The mood manipulation consisted of an autobiographical memory recall procedure conducted during the study session. After identifying positive and procedural memories suitable for use in the manipulation, participants in the Positive group recalled personally meaningful, positive, non-drug-related memories, whereas participants in the Procedural group recalled emotionally neutral autobiographical events. For each participant, research staff recorded a brief written description of each of the three autobiographical memories selected for potential use during the recall procedure and classified each as originating from childhood or another distant period of the participant’s life (“old”) versus a more recent period (“new”), to characterize the content of the memories used in the manipulation.

During the approximately one-hour MRI protocol, participants completed two 15-minute runs of a three-armed bandit reward-learning task, followed by the autobiographical memory recall mood manipulation (positive or procedural, depending on randomization), and then two additional runs of the task. Six participants (two Procedural, four Positive) who could not complete the MRI scan performed the same task sequence and memory recall procedure outside the scanner, with self-report measures administered at the same points. Self-report measures of affect, anhedonia, withdrawal symptoms, and craving were administered immediately before participants entered the MRI scanner and again immediately following completion of the scanning session. Thus, post-manipulation assessments reflected affective state following both the mood manipulation and completion of the MRI protocol. Participants also completed two additional cognitive tasks before and after the scanner, including the contextual reinforcement learning task (cRL; a probabilistic reward-learning task assessing value-based decision making) and the digit-span task (a working-memory span task). Because this manuscript is limited to self-reported affective, craving, and withdrawal outcomes, results from the neuroimaging protocol and the additional in-scanner tasks (the contextual reinforcement learning task and the digit-span task) are not analyzed here and will be reported separately. Because self-report outcomes were collected once at the start and once at the end of this approximately one-hour MRI session - rather than immediately before and after the memory recall procedure itself - the randomized comparison reported here estimates the effect of assignment to the Positive versus Procedural recall condition on end-of-session affective state, craving, and withdrawal, rather than the acute or immediate effect of the memory manipulation in isolation.

### Self-Report Measures

Affective traits and baseline clinical symptoms were assessed at baseline. Stress and arousal were evaluated using the Stress-Arousal Checklist (Mackay et al., 1978) (Stress subscale range: 0-18; Arousal subscale range: 0-12). Impulsivity and emotion regulation were assessed at baseline using the UPPS-P Negative Urgency (range: 12-48) and Positive Urgency (range: 14-56) subscales (Whiteside & Lynam, 2001; Cyders et al., 2007); these trait measures were collected to characterize the sample and are not analyzed as outcomes or predictors in the present report. Emotion regulation was also measured via the Difficulties in Emotion Regulation Scale (DERS; Gratz & Roemer, 2004; Total score range: 36-180; Impulse subscale range: 6-30). Anxiety and depressive symptoms were measured using the PROMIS Anxiety and PROMIS Depression scales (score range: 30-80) and reported as T-scores, with scores <55 considered within the normal range, 55-60 indicating mild symptoms, 60-70 indicating moderate symptoms, and >70 indicating severe symptoms. Childhood maltreatment history was assessed at screening with the 28-item Childhood Trauma Questionnaire–Short Form (CTQ; Bernstein et al., 2003; total score range: 25-125).

Positive and negative affect were assessed using the Positive and Negative Affect Schedule (PANAS; Watson et al., 1988; Serafini et al., 2016; Robinson et al., 2020), with the Positive Affect (PANAS-PA) and Negative Affect (PANAS-NA) subscales each ranging from 10 to 50. Anhedonia was assessed using the Snaith-Hamilton Pleasure Scale (SHAPS; Snaith et al., 1995; 14 items, each scored 1–4; range 14-56). Withdrawal symptoms were measured using the Amphetamine Withdrawal Questionnaire (AWQ; Srisurapanont et al., 1999; range 0-40).

Craving-related outcomes included visual digital scale (VDS) ratings of craving, desire to use stimulants, and resistance to drug use, each ranging from 0 to 10, as well as the Stimulant Craving Questionnaire (STCQ; Northrup et al., 2015; mean of 10 methamphetamine items each rated 0–6, items 10 and 19 reverse-scored; range 0–6; Cronbach’s α = .79 at both time points). For all measures, higher scores indicated greater levels of the construct assessed, except for the VDS Resist item, for which higher scores indicated greater ability to resist drug use, and the SHAPS, for which higher scores indicated greater experienced pleasure. PANAS, SHAPS, AWQ, VDS, and STCQ measures were administered immediately before and after the MRI session. The Desire for Drugs Questionnaire and Desire for Speed Questionnaire were also administered before and after the session per protocol but are not analyzed in this report, because craving is already assessed by the VDS items and the STCQ-Meth.

### Statistical Analysis

Changes in self-report outcomes from pre-to post-mood manipulation were evaluated using 2 × 2 mixed repeated-measures analyses of variance (ANOVAs), with Time (pre, post) as the within-subject factor and Group (Procedural, Positive) as the between-subject factor. Main effects of Time and Group, as well as the Group × Time interaction, were examined for each of the eight outcomes (AWQ, PANAS-PA, PANAS-NA, VDS Craving, VDS Desire, VDS Resist, STCQ-Meth, and SHAPS). Consistent with the study protocol and trial registration, PANAS-PA and PANAS-NA were the prespecified primary outcomes, and the prespecified test was the Group × Time interaction in separate 2 × 2 mixed analyses of covariance (ANCOVAs) with sex and age as covariates; the covariate-free ANOVAs are reported as sensitivity analyses. Withdrawal (AWQ) and craving (VDS items, STCQ-Meth) were named in the protocol hypothesis and are treated as secondary outcomes. SHAPS was collected before and after the session but was not named in a hypothesis and is treated as exploratory. Multiplicity was controlled within families: Holm correction across the two primary outcomes, and Benjamini– Hochberg false discovery rate (FDR) correction across the six secondary and exploratory outcomes; unadjusted and adjusted p-values are both reported. Main effects of Time, and all results for secondary and exploratory outcomes, should be interpreted as hypothesis-generating rather than confirmatory. Effect sizes were calculated using partial eta-squared (η^2^p). All statistical analyses (ANOVAs and multiple regressions) were conducted in MATLAB (The MathWorks, Natick, MA; version R2024a). Statistical significance was defined as *p* < .05 (two-tailed). Because the repeated-measures Group main effect compares groups averaged across both assessments, and the Group × Time interaction tests differential change, neither directly estimates the end-of-session difference between conditions. For each outcome, we therefore also estimated the baseline-adjusted between-group difference by regressing the post-session score on Group (Procedural = reference), the baseline score, sex, and age; this difference is reported in raw scale units (Positive minus Procedural) with its 95% confidence interval.

The regression analyses reported below address a secondary, exploratory question distinct from the primary randomized Group and Group × Time comparisons reported above; they were not part of the randomized comparison, were not specified in the study protocol or trial registration, and p-values are reported both unadjusted and FDR-adjusted across all regression coefficients. Multiple linear regression models were used to examine whether treatment group, age, abstinence duration, sex at birth, and recruitment site (Women in Recovery vs. GRAND Addiction Recovery Center) predicted pre–post change scores (post minus pre) for each affective, withdrawal, and craving-related outcome. Because each participant contributed a single pre–post change score per outcome rather than repeated within-person observations, standard (ordinary least squares) multiple regression was used rather than mixed-effects modeling. Treatment group, sex at birth, and recruitment site were entered as categorical predictors, while age and abstinence duration were entered as continuous predictors. Abstinence duration was positively skewed and was therefore compared between groups using a non-parametric Wilcoxon rank-sum test in Table 1 rather than an independent-samples t-test; it was retained in its original (untransformed) units as a continuous predictor in the regression models, consistent with standard practice for OLS regression, which assumes normality of model residuals rather than of individual predictors. Unstandardized regression coefficients (B) are reported in each outcome’s scale units per unit of the predictor (age per year, abstinence per day); categorical predictors were coded Positive = 1 (vs. Procedural), female = 1 (vs. male), and Women in Recovery = 1 (vs. GRAND). Because the ANOVA and regression models test different sets of predictors (Group and Time in the ANOVAs; Group, Age, Abstinence Duration, Sex, and Recruitment Site simultaneously in the regressions), an effect that is significant in one model but not tested in the other reflects a difference in what was modeled rather than a contradiction between analyses, and the two sets of results should not be interpreted as directly comparable tests of the same effect.

## Results

### Participant Characteristics

Across the full sample, participants demonstrated moderate baseline stress (8.92 ± 3.82) and arousal (6.18 ± 2.63), as assessed using the Stress-Arousal Checklist, moderate difficulties in emotion regulation (DERS Total: 87.93 ± 29.84), normal-to-mild anxiety symptoms (PROMIS Anxiety: 52.9 ± 9.8), and mild depressive symptoms (PROMIS Depression: 57.2 ± 10.0; **Table 1; Supplemental Figure S1**).

Groups did not differ significantly in age, sex distribution, or abstinence duration (**Table 1; Figure 1**). Groups also did not differ significantly in overall emotion-regulation difficulties (DERS Total: t_71_ = 1.96, p = .054), impulse-control difficulties (t_71_ = 0.97, p = .335), anxiety symptoms (PROMIS Anxiety: t = 0.59, p = .554), or depressive symptoms (PROMIS Depression: t = 0.53, p = .595).

**Figure 1.**
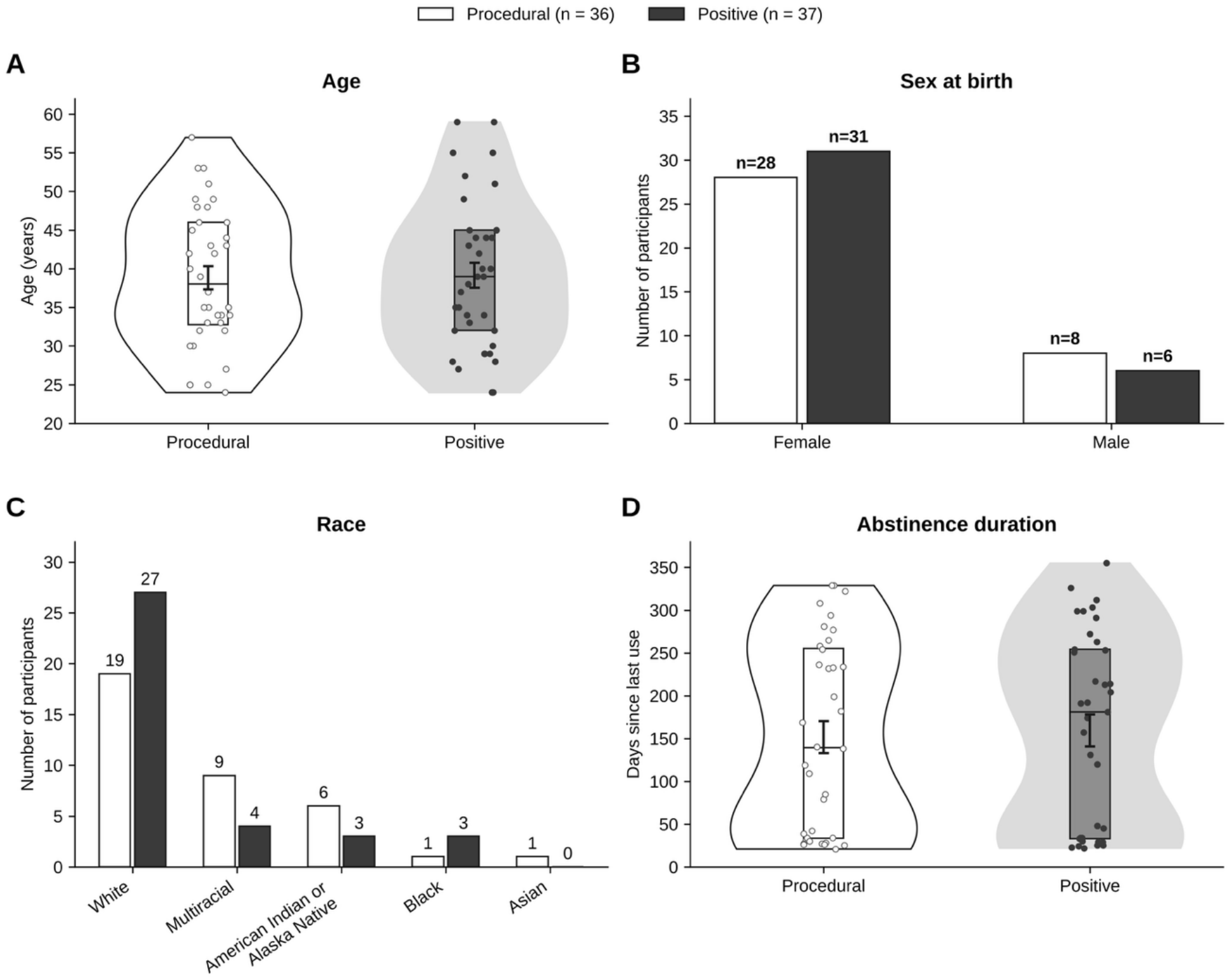
Sample Demographics. Distributions of participant **[A]** age, **[B]** sex at birth, **[C]** race, and **[D]** duration of abstinence from methamphetamine in the full sample (*N* = 73), by group (Procedural, white/open symbols; Positive, black/filled symbols). **[A]** Age (years) by group; violin plots show the distribution, boxes show the interquartile range, and error bars show mean ± SEM. **[B]** Number of participants by sex at birth (female, male). **[C]** Number of participants by race (White, Multiracial, American Indian or Alaska Native, Black or African American, Asian); race categories reflect current NIH/OMB classification standards. **[D]** Days since last methamphetamine use (abstinence duration) by group; violin plots show the distribution, boxes show the interquartile range, and error bars show mean ± SEM.

However, the Positive group reported significantly higher baseline stress and arousal than the Procedural group, as measured by the Stress-Arousal Checklist. Specifically, the Positive group had higher stress scores (9.83 ± 4.00 vs. 8.00 ± 3.44; t_70_ = −2.09, p = .041) and arousal scores (6.81 ± 2.86 vs. 5.56 ± 2.25; t_70_ = −2.06, p = .043) prior to the mood manipulation.

### Self-Report Outcomes

Table 2 summarizes results from the mixed repeated-measures ANOVAs for each outcome; detailed results are reported by outcome domain below.

**Table 2.** Mixed Repeated-Measures ANOVA of Pre-to Post-Mood Manipulation Changes in Affect, Craving, and Withdrawal Measures.

| **Outcome** | **N** | **Time (*p*)** | **Group (*p*)** | **Group × Time (*p*)** | **Group × Time, ANCOVA† p (adj. p)** | **Baseline-adjusted difference‡ [95% CI]** |
|---|---|---|---|---|---|---|
| AWQ | 73 | <0.001 | 0.331 | 0.794 | .894 (.996) | −0.21 [−1.56, 1.15] |
| PANAS Positive Affect | 73 | <0.001 | 0.022 | 0.693 | .689 (.799) | −0.53 [−2.60, 1.53] |
| PANAS Negative Affect | 73 | <0.001 | 0.772 | 0.443 | .399 (.799) | 0.54 [−0.90, 1.99] |
| VDS Craving | 73 | 0.015 | 0.755 | 0.916 | .996 (.996) | −0.02 [−0.70, 0.67] |
| VDS Desire | 73 | 0.011 | 0.791 | 0.397 | .447 (.894) | 0.31 [−0.34, 0.97] |
| VDS Resist | 73 | 0.136 | 0.673 | 0.327 | .424 (.894) | −0.38 [−2.21, 1.45] |
| STCQ-Meth | 73 | 0.405 | 0.430 | 0.612 | .617 (.925) | 0.10 [−0.15, 0.36] |
| SHAPS | 73 | 0.985 | 0.307 | 0.168 | .187 (.894) | 1.04 [−0.55, 2.63] |

**Table 3.** Multiple Linear Regression Models Predicting Pre–Post Changes in Affective Outcomes.

| **Outcome** | **N** | **Group B (p)** | **Age B (p)** | **Abstinence B (p)** | **Sex B (p)** | **Site B (p)** |
|---|---|---|---|---|---|---|
| **AWQ** | 73 | -0.03 (.962) | -0.05 (.219) | 0.009 (.135) | -1.19 (.284) | -2.03 (.185) |
| **PANAS-PA** | 73 | -0.40 (.689) | 0.03 (.541) | -0.007 (.427) | 1.07 (.503) | -0.13 (.952) |
| **PANAS-NA** | 73 | 0.77 (.426) | 0.08 (.162) | -0.002 (.824) | -2.09 (.180) | 1.31 (.540) |
| **SHAPS** | 73 | 1.01 (.207) | -0.03 (.578) | -0.001 (.869) | 0.03 (.984) | 1.56 (.378) |
| **VDS Craving** | 73 | -0.02 (.953) | 0.02 (.274) | -0.003 (.459) | 0.29 (.640) | 0.90 (.303) |
| **VDS Desire** | 73 | 0.26 (.485) | 0.02 (.342) | -0.002 (.605) | 0.13 (.836) | 0.77 (.360) |
| **VDS Resist** | 73 | -0.94 (.353) | -0.06 (.335) | -0.021 (**.021**) | -2.78 (.091) | 3.75 (.099) |
| **STCQ-meth** | 73 | 0.07 (.594) | -0.01 (.328) | 0.001 (.522) | 0.06 (.793) | -0.19 (.531) |

### Affective States

Negative affect (PANAS-NA) decreased significantly across the full sample following the mood manipulation (Time: *p* < .001), with comparable declines in both the Procedural (18.31 → 16.11) and Positive (17.51 → 16.05) groups; neither the Group main effect (*p* = .772) nor the Group × Time interaction (*p* = .443) was significant (**Table 2; Figure 2**).

**Figure 2.**
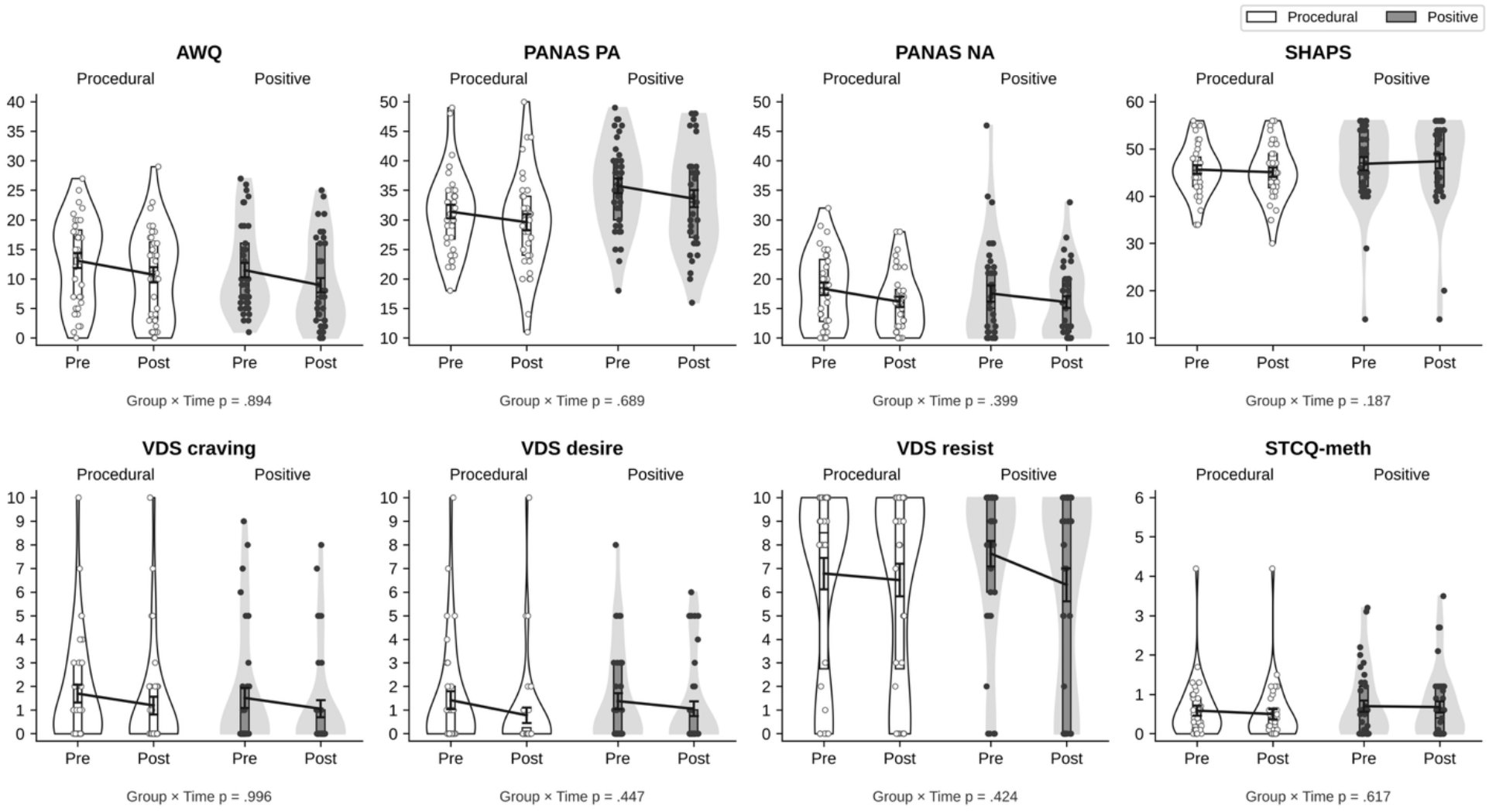
Pre-Post Changes in Affect, Withdrawal Symptoms, and Craving Following Mood Manipulation. Mean scores before and after the mood manipulation procedure are shown for the Procedural and Positive groups. Outcomes included amphetamine withdrawal symptoms (Amphetamine Withdrawal Questionnaire; AWQ), positive and negative affect (Positive and Negative Affect Schedule; PANAS-PA and PANAS-NA), visual digital scale (VDS) ratings of craving, desire to use stimulants, and resistance to drug use, stimulant craving (Stimulant Craving Questionnaire–methamphetamine version; STCQ-meth), and anhedonia (Snaith-Hamilton Pleasure Scale; SHAPS). Open circles/white violins denote the Procedural group; filled circles/gray violins denote the Positive group. Boxes show the interquartile range and median, error bars show mean ± SEM, and lines connect pre- and post-session means. The p-value below each panel is the Group × Time interaction from the prespecified mixed ANCOVA with sex and age as covariates (unadjusted; adjusted values are in Table 2).

Positive affect (PANAS-PA) also decreased significantly overall (Time: *p* < .001). Averaged across the two assessments, the Positive group reported higher PA than the Procedural group (Group: *p* = .022), but the rate of decline did not differ significantly between groups (35.73 → 33.57 vs. 31.36 → 29.58; Group × Time: *p* = .693), indicating that although the Positive group maintained higher positive affect overall (**Table 2; Figure 2**), the two groups’ trajectories from pre-to post-manipulation did not differ. Because this Group main effect averages over both assessments, including the pre-manipulation baseline, it largely reflects the groups’ baseline difference in positive affect rather than an effect of the manipulation. Adjusting for baseline PANAS-PA, sex, and age, the estimated end-of-session difference (Positive minus Procedural) was −0.53 points (95% CI [−2.60, 1.53], p = .608), providing no clear evidence of an end-of-session difference in positive affect between conditions. In the prespecified primary analysis (mixed ANCOVAs with sex and age as covariates), the Group × Time interaction was not significant for PANAS-PA (F(1, 69) = 0.16, p = .689, Holm-adjusted p = .799) or PANAS-NA (F(1, 69) = 0.72, p = .399, Holm-adjusted p = .799), and the baseline-adjusted end-of-session difference in PANAS-NA was 0.54 points (95% CI [−0.90, 1.99], p = .456). Excluding the six participants who completed the protocol outside the scanner did not change these conclusions (Group × Time: PANAS-PA p = .575; PANAS-NA p = .091; all secondary and exploratory outcomes p ≥ .13).

### Withdrawal and Craving Measures

Across the full sample, amphetamine withdrawal symptoms (AWQ; *p* < .001), craving (VDS Craving; *p* = .015), and desire to use stimulants (VDS Desire; *p* = .011) all decreased significantly from pre-to post-mood manipulation, whereas stimulant craving on the STCQ-Meth did not change significantly (p = .405). Resistance to drug use (VDS Resist) and anhedonia (SHAPS) did not change significantly over time (*p* = .136 and *p* = .985, respectively). No significant Group × Time interactions were observed for any outcome (all *ps* ≥ .168), indicating that pre-to post-session changes did not differ between the Positive and Procedural groups (**Table 2; Figure 2**). There were also no significant main effects of Group for these outcomes (all *ps* ≥ .306). Because the two reverse-worded STCQ-Meth items had low item–total correlations (r = .00–.42; Cronbach’s α = .79 for all 10 items and .87–.90 for the eight forward-worded items), we repeated the STCQ-Meth analysis using only the eight forward-worded items. This score decreased over the session (p = .015), but the Group × Time interaction remained non-significant (p = .472).

### Predictors of Affective Change

Treatment group and age were not significantly associated with pre–post changes in any affective, withdrawal, or craving-related outcome (all *p*s > .05). Neither sex at birth nor recruitment site was significantly associated with pre–post change in any outcome (all ps ≥ .09). Because all 14 male participants were recruited from GRAND Addiction Recovery Center, sex and site are confounded in these models (variance inflation factors 1.6 and 4.7, respectively). Abstinence duration significantly predicted change in resistance to drug use (VDS Resist; B = −0.02 points per day, *p* = .021), such that longer abstinence duration at study entry was associated with a steeper decline in resistance to drug use from pre-to post-session. No other significant associations were observed. After Benjamini–Hochberg FDR correction across all 40 regression coefficients, this association did not remain significant (FDR-adjusted p = .801).

## Discussion

The present study examined whether a brief positive autobiographical memory recall manipulation could differentially improve affective state, withdrawal symptoms, and craving among treatment-seeking individuals with MUD, relative to a neutral, procedural memory recall condition. Contrary to our primary hypothesis, the Positive and Procedural groups did not differ in their pattern of change for withdrawal symptoms or craving-related outcomes, and the Group × Time interaction was non-significant for every outcome examined, including the prespecified primary outcomes of positive and negative affect. Instead, a robust main effect of Time emerged across the full sample: participants demonstrated significant reductions in withdrawal symptoms, negative affect, and craving over the course of the session regardless of group assignment.

Positive affect was the only outcome with a main effect of Group in the repeated-measures ANOVA; because that effect averages across both assessments, it largely reflects the groups’ baseline difference, and after adjustment for baseline PANAS-PA, sex, and age the end-of-session difference was small and imprecise (−0.53 points, 95% CI [−2.60, 1.53]). Taken together, no outcome showed a robust randomized-group effect once baseline differences and other covariates were accounted for. Overall, although affect, craving, and withdrawal changed across the session in both groups, we found no clear evidence of an end-of-session difference between the positive and procedural recall conditions.

Methodological and timing constraints inherent to embedding the manipulation within a neuroimaging protocol offer a primary explanation for these findings. Post-manipulation self-reports were administered only after completion of the entire MRI protocol - approximately one hour after baseline - rather than immediately following the memory recall procedure. While positive autobiographical recall can produce immediate mood elevation (Westermann et al., 1996), transient affective shifts likely decayed during subsequent task performance and extended scanner exposure (Gilet, 2008; Joseph et al., 2020). Furthermore, prolonged confinement, background noise, and sustained attention demands are known to induce fatigue (Mrazek et al., 2016; Tessitore et al., 2002), selectively depleting high-arousal positive affect without necessarily elevating threat- or distress-related negative affect (Inzlicht et al., 2014; Watson & Tellegen, 1985). This resource depletion likely accounts for the overall decline in positive affect and may have obscured condition-specific mood improvements. To isolate acute memory-driven effects from scanner-induced decay, future studies should capture brief, interval-based self-reports immediately before and after the memory recall procedure prior to subsequent scanning protocols.

The qualitative characteristics of the recalled memories may have further attenuated the manipulation’s intended effects. Although participants in the positive condition were instructed to select non-drug-related memories, individuals with extensive substance use histories frequently drew upon childhood or otherwise distant (“old”) recollections: research staff classified at least one of the three candidate memories as an “old” memory for 92% of Positive-group participants, compared with only 6% of Procedural-group participants. While objectively positive, recalling life circumstances from before the onset of addiction can trigger bittersweet emotional states, grief, regret, and nostalgia (Sedikides & Wildschut, 2016). Consistent with this, prior work suggests the relationship between positive memory and substance use is not uniform: among trauma-exposed individuals, higher counts of retrieved positive memories were associated with greater hazardous substance use specifically among those with difficulties regulating positive emotion (Compton et al., 2023), suggesting that individual differences in tolerating and regulating positive states may have shaped participants’ responses to the memory manipulation here as well. Because autobiographical memory retrieval frequently elicits complex, mixed emotional responses (Larsen et al., 2001; Larsen & McGraw, 2011), these “contaminated” recollections may have blunted pure positive affect elevation. Additionally, participants reported only mild baseline depressive symptoms on average, potentially creating a ceiling effect that limited measurable improvement. Future work should incorporate standardized pre-screening to quantify memory age, nostalgia, and emotional complexity to prevent mixed-emotion contamination.

Despite the lack of between-group divergence, participants in both conditions showed significant reductions in craving, withdrawal, and negative affect from pre-to post-session. Because both conditions completed the same sequence of memory recall, cognitive tasks, MRI scanning, and repeated self-report assessments, this design cannot distinguish which feature of the study visit, if any, produced these session-level changes. Structured cognitive engagement, such as the reward-learning tasks completed during scanning, could plausibly divert attention away from withdrawal-related and drug-cue processing (Goldstein & Volkow, 2011; May et al., 2011; Parvaz et al., 2012; Van Dillen & Andrade, 2016); alternatively, active, purposeful engagement in either recall task, consistent with behavioral activation models, or simply participating in a compensated study visit, could also plausibly contribute to mood and symptom change independent of task content (Daughters et al., 2008, 2018; Mazzucchelli et al., 2009; Ryan & Deci, 2000). Because the present design does not distinguish among these candidate explanations, we describe the whole-group changes as session-level observations rather than evidence for a specific mechanism, and we do not interpret them as clinical improvement or as evidence of reduced relapse risk. A passive rest control condition and explicit manipulation of agency in future designs would be needed to test these candidate mechanisms directly.

This study also has several sample-level limitations. The Procedural and Positive groups differed at baseline in comorbid psychiatric diagnoses: participants assigned to the Procedural condition were more likely to meet criteria for PTSD than those assigned to the Positive condition (34.4% vs. 8.6%; Fisher’s exact p = .015; Table 1). Because PTSD is associated with altered autobiographical memory processing and affective reactivity, this baseline imbalance is a potential confound that the present sample size did not allow us to formally test or statistically adjust for, and it should be considered when interpreting the group comparisons reported above.

## Acknowledgements and Disclosures

MCMG acknowledges support from NIGMS (Grant number: P20GM121312; PI: M.P. Paulus). The findings and conclusions of this article are those of the authors and do not necessarily represent the views and opinions of these organizations, which had no role in designing or planning the study, in collecting, analyzing, or interpreting the data, in writing the article, or in deciding to submit it for publication.

The study was approved by the WCG Institutional Review Board (LIBR protocol #2024-002), all participants provided written informed consent before participation, and the trial is registered at ClinicalTrials.gov (NCT06410196). Data are available from the corresponding author (MCMG) on reasonable request.

These findings were presented by SIQ in poster format at the Oklahoma Psychological Association conference in Tulsa, OK, in November 2025.

SIQ collected the data. SIQ and MCMG accessed the data, performed data cleaning and data analysis, and contributed to result interpretation. SIQ wrote the first draft of the article and edited subsequent drafts. MCMG designed the study, guided and oversaw the data analysis, helped with result interpretation, edited drafts of the article, and provided supervision and guidance. JLS, MI, MPP, and MCMG reviewed and edited subsequent drafts of the manuscript. All authors made substantive contributions to the article, read, and approved the final article.

The authors report no biomedical financial interest or potential conflicts of interest.

