## Supplemental for "Influence of Memory Modulation on Affective States in Methamphetamine Use Disorder"

**Figure S1. Baseline Clinical Characteristics**

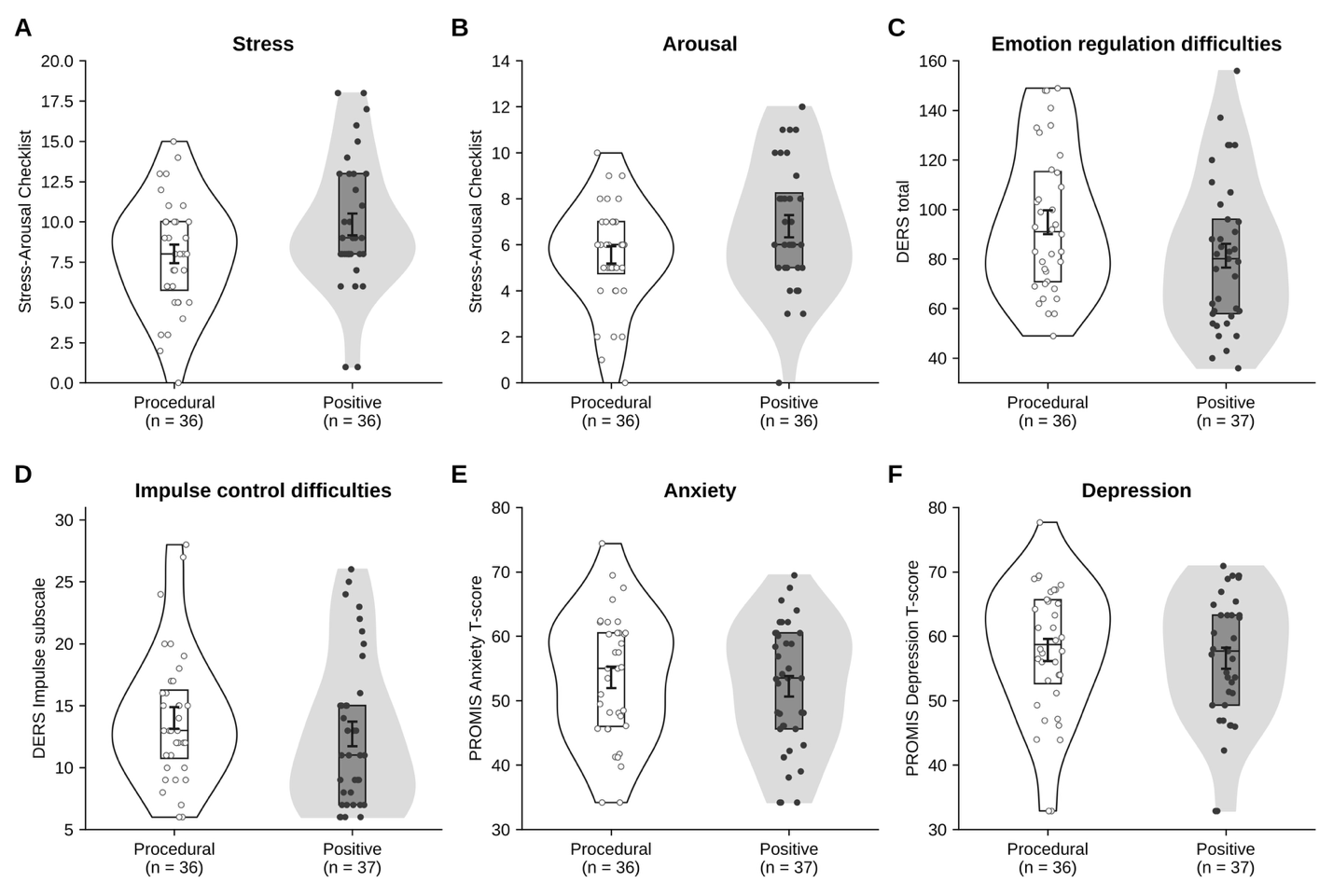

***Figure S1.*** Baseline (A) stress and (B) arousal (Stress-Arousal Checklist), (C) emotion regulation difficulties (DERS total), (D) impulse control difficulties (DERS Impulse subscale), and (E) anxiety and (F) depression (PROMIS T-scores) for the Procedural (white) and Positive (gray) groups. Violins show the distribution, boxes show the interquartile range and median, and error bars show mean ± SEM; the number of participants with data is shown under each panel. UPPS-P urgency scores were available for only seven participants and are not shown.

**Figure S2. Participant Flow**

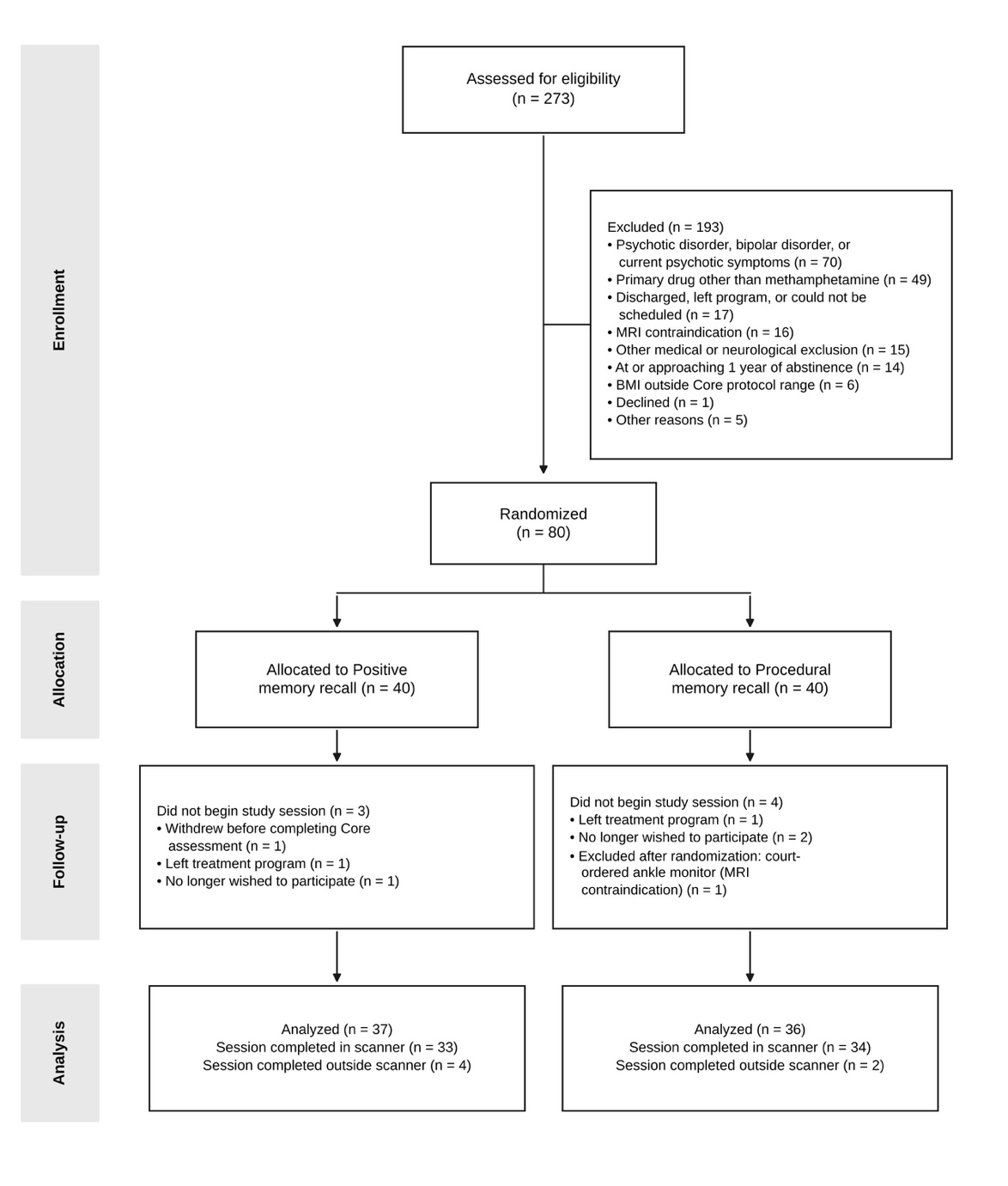

***Figure S2.*** Flow of participants through screening, randomization, and analysis (CONSORT). Exclusion reasons were recorded as free text in the study tracker and grouped into the categories shown, with the first-listed reason taking priority. None of the seven participants lost after randomization completed the pre-session assessment. Six analyzed participants completed the same task sequence and memory recall procedure outside the scanner.
